# Step-wise evolution of temperature-mediated phenotypic plasticity in eyespot size across nymphalid butterflies

**DOI:** 10.1101/378836

**Authors:** Shivam Bhardwaj, Lim Si-Hui Jolander, Markus R. Wenk, Jeffrey C. Oliver, H. Frederik Nijhout, Antónia Monteiro

## Abstract

There are two disparate views regarding phenotypic plasticity. One regards plasticity as a derived adaptation to help organisms survive in variable environments^1, 2^ while the other views plasticity as the outcome of flexible, non-canalized, developmental processes, ancestrally present in most organisms, that helps them colonize or adapt to novel environments^3–5^ e.g., a pre-adaptation. Both views of plasticity currently lack a rigorous, mechanistic examination of ancestral and derived states and direction of change^2^. Here we show that the origin of phenotypic plasticity in eyespot size in response to environmental temperature observed in *Bicyclus anynana* butterflies is a derived adaptation of this lineage. Eyespot size is regulated by temperature-mediated changes in levels of a steroid hormone, 20E, that affects proliferation of eyespot central cells expressing the 20E receptor (EcR)^6, 7^. By estimating the origin of the known physiological and molecular components of eyespot size plasticity in a comparative framework, we showed that 20E titer plasticity in response to temperature is a pre-adaptation shared by all butterfly species examined, whereas the origin of expression of EcR in eyespot centers, and the origin of eyespot sensitivity to the hormone-receptor complex are both derived traits found only in a subset of species with eyespots. The presence of all three molecular components required to produce a plastic response is only observed in *B. anynana*. This gradual, step-wise, physiological/molecular response to temperature is a likely adaptation to temperature variation experienced across wet and dry seasons in the habitat of this species. This work supports, thus, the first view of plasticity as a derived adaptation.

---

The two views on phenotypic plasticity articulated above, as an adaptation or a pre-adaptation, require either that plasticity evolves under natural selection or that it is ancestral and widespread and facilitates adaptation. Several case studies have been documented in support of the first^8–10^, and second evolutionary scenarios^11, 12^ but to date, almost nothing is known about how the original plastic responses underlying both hypotheses originated and evolved at the proximate, mechanistic level. Details of how plasticity originates, and whether or not it is widespread and ancestral to a group of species, regardless of their current living environments, may also help discriminate between plasticity being a facilitator or a consequence of organismal adaptation.

A comparative approach that addresses the mechanistic origins of plasticity needs grounding in a sufficiently well understood molecular mechanism of plasticity. Here we use dramatic seasonal variation in the size of *B. anynana* wing eyespot patterns as our case study. *Bicyclus* species live throughout dry and wet seasons in Africa, where eyespots of different sizes serve different ecological roles^13, 14^. In the hot wet season, the large exposed ventral eyespots help deflect attacks of invertebrate predators towards the wing margins^15^, whereas in the cool dry season the smaller eyespots help in camouflage against vertebrate predation^16^.

Eyespot size plasticity in *B. anynana* is mostly controlled by temperature, which leads to variable titers of the hormone 20-hydroxyecdysone (20E) at the wandering (Wr) stage of larval development^6^. Manipulations of 20E signaling alone, at that time in development, are sufficient to modify eyespot size^6^. Upon sufficient 20E signaling, these central cells divide and produce a larger central group of signaling cells^7^ and ultimately a larger eyespot. Given knowledge of how eyespot size plasticity functions in one species, we sought to investigate how this system of temperature sensitivity evolved by performing a comparative study across nymphalid butterflies, with and without eyespots.

Eyespots originated once within the nymphalid family, about 85 mya, likely from pre-existing simple spots of a single colour^17, 18^ but it is unclear whether size plasticity in response to temperature evolved before or after the origin of eyespots. If eyespot or spot size plasticity is an ancestral pre-adaptation, it is possible that even species of butterflies that do not experience seasonal environments (such as those living near the equator), might have the ability to develop different eyespot or spot sizes when reared at different temperatures under experimental conditions. Alternatively, if eyespot size plasticity is an evolved adaptation, used exclusively by species living in seasonal environments, then only these species should exhibit plasticity.

To test these hypotheses, we reared twelve species from different nymphalid sub-families, and from tropical, or sub-tropical regions, plus one outgroup papilionid species (Table S1) at two different temperatures, separated by 10 degrees Celsius, and measured eyespot size plasticity in adult females. Three different types of reaction norm to rearing temperature were observed across species (Fig. 1A). Five species showed no significant difference in hindwing (HW) Cu1 eyespot size when reared across two temperatures and were deemed not plastic. Most species showed a decrease in eyespot size with an increase in temperature and had a negative slope in their reaction norms. *B. anynana* was the only species which displayed a positive slope in its reaction norm, where eyespot size increased with temperature^13^ (Table S2). Ancestral character state reconstructions for the slope of these reaction norms suggested that eyespot size plasticity of any form is a derived trait within nymphalids, with three or four possible independent origins. Ancestral species of nymphalids lacked plasticity, whereas there were two or three independent origins of a negative response of eyespot size to increasing temperature and a separate origin of the opposing pattern of plasticity in ventral HW eyespot size in the lineage leading to *B. anynana* (Fig. 1B).

**Figure 1.**
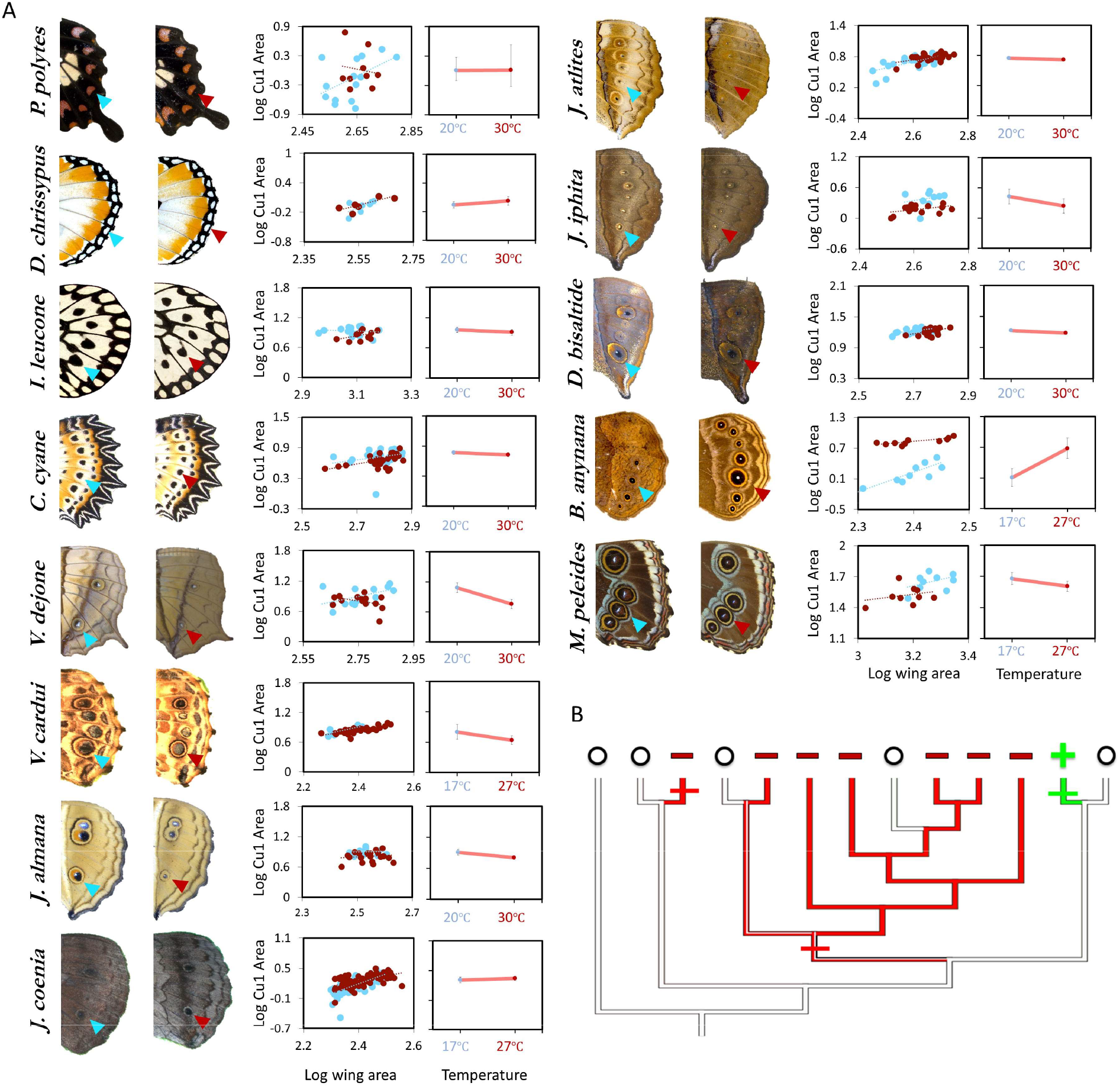
Eyespot/spot size plasticity is widespread across butterfly lineages but the response to rearing temperature has different norms of reaction across species. A. Size of hindwing ventral Cu1 eyespots (arrowheads). Thirteen species of butterflies were reared at two different rearing temperatures. Eyespot size corrected for wing size is plotted for two different temperatures (low temperature 17°C or 20°C is marked with blue symbols, while high temperature of 27°C or 30°C is marked with red symbols). Error bars represent 95% CI of means. B. Phylogenetic analysis suggests 3 independent origins for two different patterns of plasticity (eyespot size decreases with increasing temperatures: red lineages, and eyespot size increases with increasing temperature: green lineage). The ancestral reconstruction for the gain of negative plasticity is equivocal for two (shown) or three (not shown) gains. That is, it is equally parsimonious that negative plasticity was gained as shown or that it was gained three separate times: once leading to *I. leucone*, once leading to *V. dejone*, and once leading to the MRCA of *V. cardui* and *D. bisaltide*.

To investigate the molecular basis for these different patterns of plasticity we looked at 20E titers and EcR expression across species using female data. 20E titers at the Wr stage were consistently higher at the higher rearing temperature across all butterflies (Fig. 2a) (Table S3), suggesting that 20E titer plasticity in response to temperature is an ancestral trait shared across these butterflies. EcR expression at the Wr stage was absent from spot centers in species with simple spots, but was present in the eyespot central cells across all species investigated, with a few exceptions (*Junonia coenia and Junonia almana*) (Fig. 2B). This suggests that EcR localization in eyespots is a derived trait, present only in species with eyespots.

**Figure 2.**
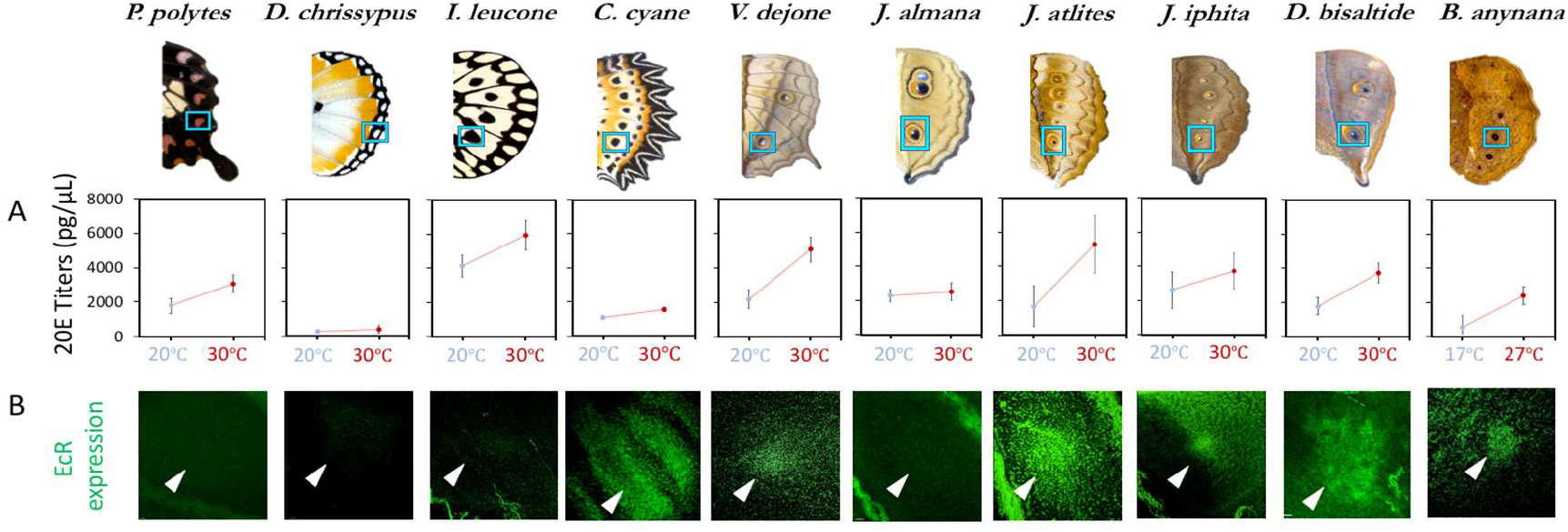
20E titers increase with rearing temperature across most species but EcR expression in eyespots is only found in a subset of nymphalids. **A.** 20E titers increase with an increase in rearing temperature across most species. This trait is ancestral in nature, with a likely origin before the origin of eyespots. **B.** EcR is absent in simple spots, but present in the future eyespot centers of most of the species investigated (N ≥ 3 for each immunostaining).

Finally, to test whether eyespots expressing EcR are size regulated by 20E we manipulated 20E levels and EcR receptors directly. Functional experiments were performed in four species of butterflies from different Nymphalid subfamilies, *Idea leuconoe* (Danainae), a control outgroup danainae with no EcR expression in its black spots, *Vindula dejone* (Nymphalinae), *Doleschallia bisaltide* (Nymphalinae), and *B. anynana* (Satyrinae), the latter three displaying EcR expression in their eyespot centers. Our prediction would be that *Idea* should not respond to 20E signaling at all, given the lack of the receptor in its spots, and that increases in 20E signaling at low temperature might cause the eyespots of *Vindula* and *Doleschalia* to become smaller but those of *B. anynana* to become larger, whereas decreases of 20E signaling at high temperature might cause the eyespots of the first two species to become larger but smaler in *B. anynana*. Injections of 20E into female wanderers reared at low temperature (and with lower 20E titers) and of an EcR antagonist, CucB, into female wanderers reared at high temperature (and with higher 20E titers), showed no response across the first three species, whereas eyespot size significantly increased with 20E injections and decreased with antagonist injections in *B. anynana* (Fig. 3). These data indicate that only the eyespots of *B. anynana* are sensitive to 20E signaling, within the natural range of titers displayed by these species. This sensitivity is a derived trait potentially restricted to the satyrid sub-family within nymphalids (Fig. 4).

**Figure 3.**
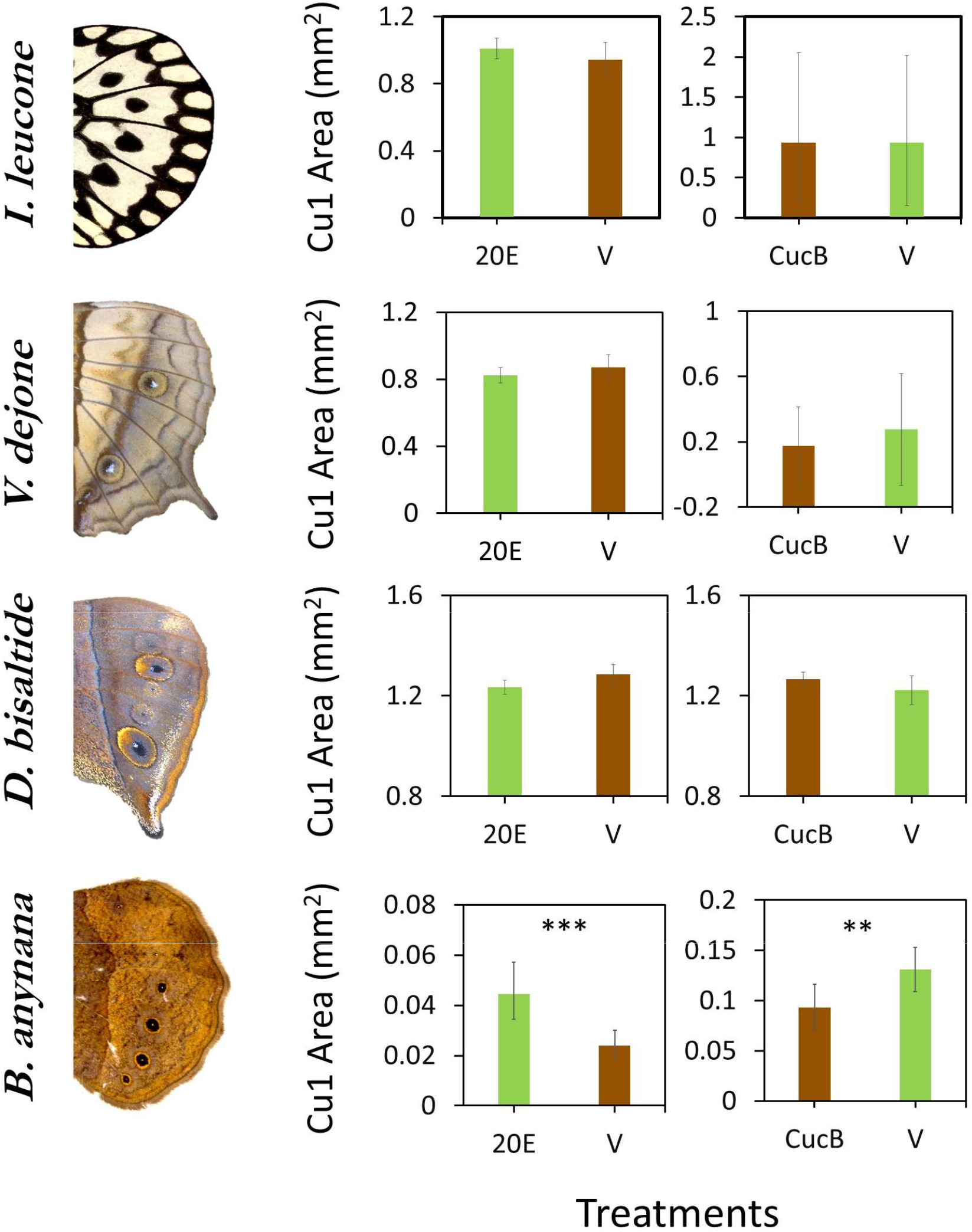
Sensitivity of eyespots to EcR mediated signaling originated in the lineage leading to *B. anynana* butterflies. Four species of butterflies were injected with 20E hormones or EcR antagonists (CucB) during the Wr stage. While *Idea leuconoe, Vindula dejone and Doleschallia bisaltide* are not sensitive to either of the hormone signal manipulations, *B. anynana* shows sensitivity towards both 20E and CucB. Error bars represent 95% CI of means. Significant differences between treatments are represented by asterisks: **, p < 0.01, ***, p < 0.001.

**Figure 4.**
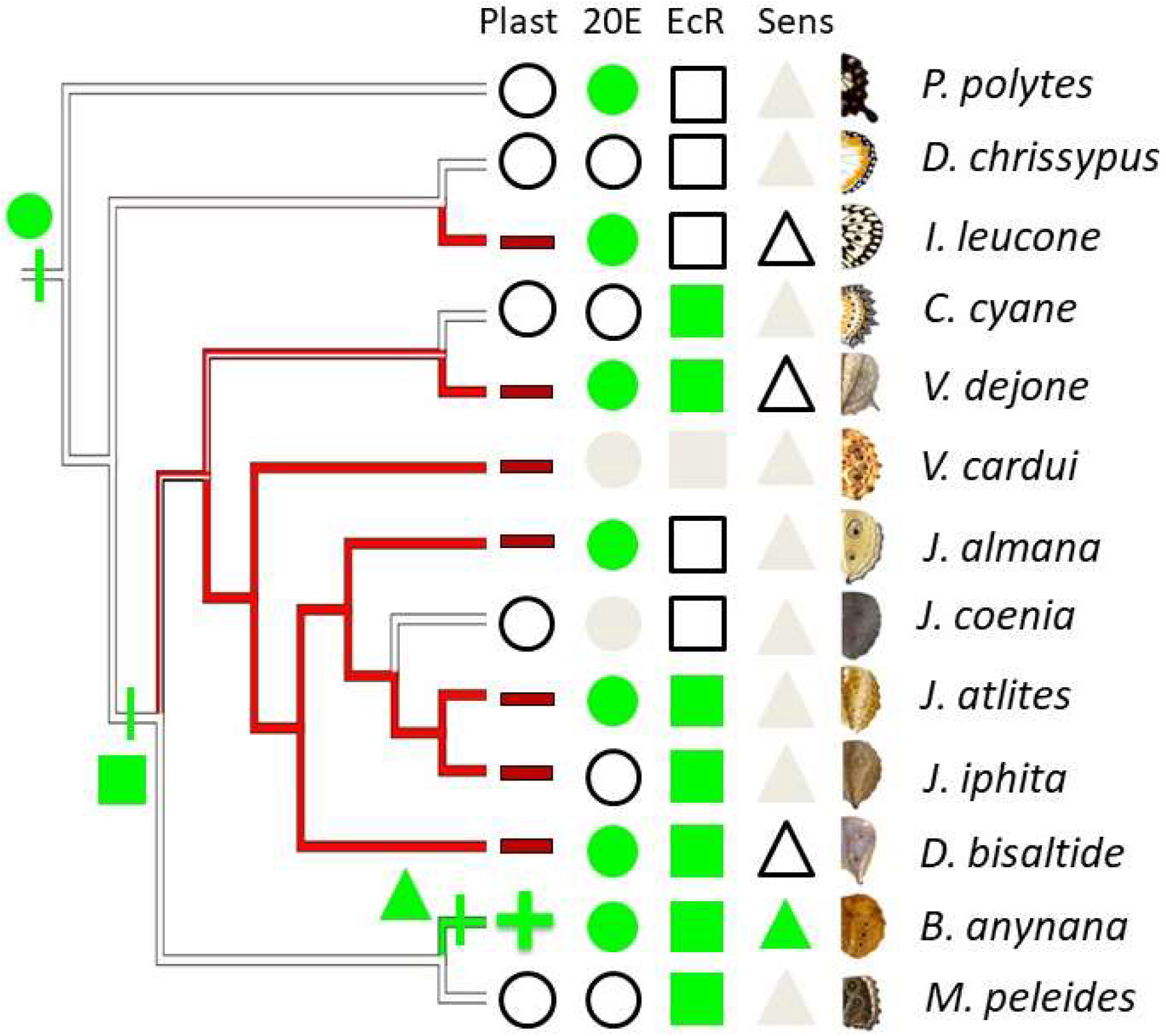
Phenotypic plasticity as a complex trait originated gradually. Phylogenetic analysis suggests three or four independent origins for two different patterns of plasticity (eyespot size decreases with increasing temperatures: red lineages and - sign, and eyespot size increases with increasing temperature: green lineage and + sign). Green circles (character state 1) represents high 20E titers with increasing temperature, while white circles (character state 0) represent no significant difference in titers at two developmental temperatures. Green squares represent presence of EcR in eyespots, while white squares represent its absence. EcR expression in eyespots is inferred to have originated concurrently with the origin of eyespots, about 85 Mya, and subsequently lost in a few nymphalid lineages. Green triangles represent sensitivity towards 20E in respective species (character state 1), while white triangles represent absence of sensitivity (character state 0). Gray circles, squares and triangles represent missing data points. Alternative models using Maximum Likelihood reach similar conclusions (Supplementary Information).

While multiple reports have focused on the role of hormones as mediators of trait plasticity^19–22^, the physiological and developmental details of how a fully functional plastic trait originates during the course of evolution were still obscure. Here we identified the approximate evolutionary origins of individual components of a plastic response of eyespot size in response to temperature and discovered this plastic response to be a complex trait that evolved gradually via changes to different molecular components. Our work showed that the origin of plasticity in hormone titers, the origin of hormone receptor expression in the trait, and the origin of eyespot sensitivity to these hormones all took place at different stages of nymphalid diversification (Fig. 4).

An increase in eyespot size in response to temperature appears to be restricted to satyrid butterflies, and is a derived response within nymphalids. Plasticity in eyespot size in butterflies had been primarily documented in satyrid butterflies such as *Melanitis leda* and several *Bicyclus* species^23–25^ where size was always found to positively increase with rearing temperature. Most of the reared species of nymphalids and the papilionid species showed a slight decrease in eyespot/spot size with an increase in temperature, while some species showed no plasticity at all. This decrease in eyespot size with increasing temperature may simply reflect non-adaptive variation from a poorly canalized system. In addition, satyrid butterflies, but none of the other species, used the 20E asymmetry to regulate the size of their eyespots in a novel way. This was enabled by the prior recruitment of EcR to the eyespot central cells perhaps concurrently with eyespot origins (Fig. 4). These central signaling cells play an important role in determining eyespot size at the Wr stages of development^26^. Some species, such as *Junonia coenia*, retain expression of EcR in eyespots but only at other stages of wing development^27^. Finally, the active 20E-EcR complexes increase eyespot size in *B. anynana* but not in other species with similar EcR expression in their eyespot centers. The ability of 20E to promote localized patterns of cell division might have evolved in the lineage leading to *B. anynana* alone^7^.

Eyespot size plasticity in connection with wet and dry seasonal forms is widely conserved across the sub-family Nymphalinae^28^ but our results suggest that different mechanisms may have evolved to regulate eyespot size plasticity in these lineages. Our controlled rearing experiments showed that all nymphalinae (*Vanessa cardui, Junonia almana, J. coenia, J. atlites, J. iphita* and *Doleschallia bisaltide*) produced only small changes in the size of the their Cu1 eyespots in response to rearing temperature, and these were in the opposite direction to those observed in *B. anynana*. Other environmental factors might cue and regulate these species’ seasonal morphs (Fig. S2), perhaps cues that better predict the arrival of the seasons where these butterflies have evolved. Investigations at the proximate level will be required to correctly establish the environmental cues that induce seasonal forms in these other butterfly species. For now, we uncover phenotypic plasticity in eyespot size in *B. anynana* as a complex, step-wise adaptation to seasonal environments cued by temperature that required very specific mutations to originate. This work also serves as a warning that if all forms of plasticity are as specific and hard to evolve as the one documented in *B. anynana*, these exquisite adaptations to specific predictable fluctuating environments may in fact, lend the species vulnerable to extinction under unpredictable climate change, as previously noted ^29^.

## Supplementary Information

### Materials and methods

#### Butterfly husbandry

*B. anynana* were raised from lab populations in Singapore, under temperature control chambers at 17°C and 27°C, with a 12h light: dark cycle and 80% RH. *Vanessa cardui*, and *Morpho peleides* were reared in climate chambers at Yale University, New Haven at 17°C and 27°C. *Junonia coenia* was reared at 20° and 30°C at 16H:8H light: dark cycle at Duke University. All other species of butterflies were reared at Entopia (formerly, Penang Butterfly Farm, Penang, Malaysia) in temperature controlled chambers (PT2499 Incubator, Exoreptiles, Malaysia) at 20°C and 30°C. 70% RH was maintained and monitored using (PT2470 Hygrometer, Exoreptiles, Malaysia) and EL-USB-2 data loggers (Lascar Electronics, PA 16505, USA).

Four hours after emergence, butterflies were captured and frozen in glassine envelopes at −20°C. All larvae in this experiment were sexed during larval or pupal stages and only females were used for analysis. Wings were carefully dissected and imaged using a Leica upright microscope. Wing images were processed in ImageJ, where area and eyespot size were measured using selection tools.

#### Haemolymph collection

Previous studies in *B. anynana* have pointed to the wandering (Wr) stage as the critical temperature sensitive stage for determination of ventral hindwing eyespot size^6^. Time lapse photographs of larval development were captured using a RICOH camera to determine the beginning of the Wr stage across all species. Initiation of Wr stage is marked by the larvae stopping to feed, purging their gut, and starting to wander away from the food and looking for a place to pupate. Using Hamilton syringes, 20uL of haemolymph, were extracted from each larvae at ~70% development in Wr stage (15h after Wr started for animals reared at 30°C, and 25h for animals reared at 20°C). Extracted haemolymph was then dissolved in freshly prepared 90ul methanol + 90 ul isooctane and stored at −20°C until hormone extraction^7^.

#### Wing tissue collection

Larval wing discs were dissected from Wr stage larvae and stored in fix buffer until further processing at 4°C. These were later stained for EcR expression using a primary antibody 15F1 (DSHB) raised against a *Manduca sexta* EcR peptide shared across all isoforms of EcR, and secondary antibody AlexaFlour 488 green. Spalt, a nuclear marker for spots and eyespots, was used as a location marker for putative eyespots/spots in the larval wings. Serial optical sections of the Cu1 eyespot wing sector were imaged using LSM510 Meta, to distinguish between dorsal and ventral surfaces. Specific slices were obtained from raw images using Imaris v8.64 (ImarisXT, Bitplane AG, software available at http://bitplane.com. *Junonia coenia* EcR data were taken from Koch and Nijhout, 2003^27^.

#### 20E and antagonist injections

Four species of butterflies, *Idea leuconoe, Vindula dejone, Doleschallia bisaltide*, and *B. anynana*, were injected with 20E or CucB during the Wr stage. Injections were made at ~50% development of Wr stage (12-14h at 30°C, 18-22h at 20°C; For *B.anynana*, rearing were done at 27°C and 17°C respectively). Average body weights of wandering larvae and total haemolymph present were calculated for each species, and used to calculate naturally circulating 20E levels *in vivo*. A gradient of different concentrations of 20E and CucB were used for pilot experiments. Maximum concentrations of 20E, which did not surpass the natural levels, and of CucB, which did not cause mortality or pupation defects, were used for injections and are summarized in the table below. 20E and CucB were dissolved in 10% EtOH to make working solution for injections. Equal volume injections of Vehicle (10% EtOH in Saline) injections were done as controls. After injections, animals were reared at their regular rearing temperature (17°C for *B.anynana*, 20°C for other 20E injected animals and 27°C for *B.anynana*, 30°C for CucB injected animals) until emergence as adults. After emergence, the wings were dissected, imaged, and scored for further analysis.

#### Statistical analysis

All wing and eyespot data were log10 transformed to ensure linearity of wing size with eyespot size for purposes of allometric scaling and regression analysis, and to be able to compare slopes across species with different eyespot sizes and wing sizes. Univariate ANCOVAs were performed using hindwing Cu1 eyespot area as the main variable, hindwing area as a covariate, and rearing temperature as a fixed factor in SPSS v21. Graphs were plotted in Microsoft Office 2016 for Mac. Slopes for plasticity of eyespot size and 20E titers were measured using the expression:

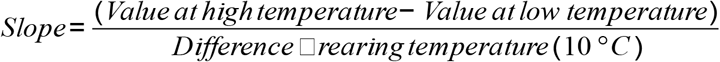

Using reverse transformed data for eyespot size, and untreated values for hormone titers.

**Table S1:** Species reared for comparative morphometrics, gene expression and hormonal measurements

| Species | Family/Nymphalid Subfamily | Spots/Eyespots | Rearing temp. (°C) | Source of animals used in experiments | Climatic conditions (Köppen classification) | Reported Seasonality in Spot/Eyespot Size |
| --- | --- | --- | --- | --- | --- | --- |
| *Junonia atlites* | **Nymphalidae**/Nymphalinae | Eyespots | 20/30 | Malaysia | Tropical; Equatorial humid (Af) | No |
| *Junonia coenia* | **Nymphalidae**/Nymphalinae | Eyespots | 20/30 | USA | Subtropical | No |
| *Junonia iphita* | **Nymphalidae**/Nymphalinae | Eyespots | 20/30 | Malaysia | Tropical; Equatorial humid (Af) | No |
| *Junonia almana* | **Nymphalidae**/Nymphalinae | Eyespots | 20/30 | Malaysia | Tropical; Equatorial humid (Af) | Yes |
| *Doleschallia bisaltide* | **Nymphalidae**/Nymphalinae | Eyespots | 20/30 | Malaysia | Tropical; Equatorial humid (Af) | No |
| *Vanessa cardui* | **Nymphalidae**/Nymphalinae | Eyespots | 17/27 | USA | Subtropical | No |
| *Vindula dejone* | **Nymphalidae**/Heliconinae | Eyespots | 20/30 | Malaysia | Tropical; Equatorial humid (Af) | No |
| *Cethosia cyne* | **Nymphalidae**/Heliconinae | Eyespots | 20/30 | Singapore, Malaysia | Tropical; Equatorial humid (Af) | No |
| *Bicyclus anynana* | **Nymphalidae**/Satyrinae | Eyespots | 17/27 | Africa | Tropical; Equatorial, winter dry (Aw) | Yes |
| *Morpho peleides* | **Nymphalidae**/Morphinae | Eyespots | 17/27 | Costa Rica | Subtropical | No |
| *Danaus chrysippus* | **Nymphalidae**/**Danainae** | Spots | 20/30 | Malaysia | Tropical; Equatorial humid (Af) | No |
| *Idea leuconoe* | **Nymphalidae**/**Danainae** | Spots | 20/30 | Taiwan | Subtropical; Warm, humid, hot summers <sup>30</sup> | No |
| *Papilio polytes* | **Papilionidae - Outgroup** | Spots | 20/30 | Malaysia | Tropical; Equatorial humid (Af) | No |

**Table S2:** F statistics, p-values from analysis of covariance for differences in Cu1 eyespot size between rearing temperatures (fixed factor) and assigned character state for phylogenetic analysis. Wing size was used as a covariate. Rows highlighted in red indicate species where eyespot size decreases significantly with rearing temperature (negative slope). Species highlighted in green shows the opposite pattern (a significant positive slope). Character states of −1 = negative slope; 0=no plasticity; 1=positive slope.

| Species | Family/<br>Subfamily | Factor | F stats | P value | DF (Factor,<br>Error) | Slope of<br>reaction norm | Character<br>state |
| --- | --- | --- | --- | --- | --- | --- | --- |
| *Papilio polytes* | **Papilionidae** | temp. | 0.360 | 0.360 | 1,59 | 0.000 | 0 |
| *Danaus chrysippus* | **Danainae** | temp. | 0.318 | 0.585 | 1,59 | 0.008 | 0 |
| *Idea leucone* | **Danainae** | temp. | 10.073 | 0.031 | 1,59 | -0.004 | -1 |
| *Cethosia cyane* | Heliconinae | temp. | 0 | 0.096 | 1,59 | -0.004 | 0 |
| *Vindula dejone* | Heliconinae | temp. | 8.247 | 0.009 | 1,59 | -0.032 | -1 |
| *Vanessa cardui* | Nymphalinae | temp. | 15.056 | 0.001 | 1,59 | -0.016 | -1 |
| *Junonia almana* | Nymphalinae | temp. | 15.832 | <0.001 | 1,59 | -0.010 | -1 |
| *Junonia coenia* | Nymphalinae | temp. | 0.888 | 0.352 | 1,59 | 0.003 | 0 |
| *Junonia atlites* | Nymphalinae | temp. | 4.683 | 0.011 | 1,59 | -0.002 | -1 |
| *Junonia iphita* | Nymphalinae | temp. | 11.670 | 0.042 | 1,59 | -0.018 | -1 |
| *Doleschallia bisaltide* | Nymphalinae | temp. | 13.170 | 0.001 | 1,59 | -0.005 | -1 |
| *Bicyclus anynana* | Satyrinae | temp. | 42.769 | <0.001 | 1,59 | 0.057 | 1 |
| *Morpho peleides* | Morphinae | temp. | 0.765 | 0.393 | 1,19 | -0.007 | 0 |

**Table S3:** F statistics, p-values from analysis of covariance for differences in 20E hormone titers between rearing temperatures (fixed factor) and assigned character states for phylogenetic analysis. Wing size was used as a covariate. All data were log10 transformed to ensure linear allometries and comparable variances across temperature treatments. Rows highlighted in green indicate species where 20E titers increase significantly with rearing temperature (positive slope). Character state of 0=no plasticity; 1=positive slope.

| Species | Factor | F stats | Slope of reaction norm | P value | DF (Factor, Error) | Character state |
| --- | --- | --- | --- | --- | --- | --- |
| *Papilio polytes* | Temperature | 0.004 | 126.087 | <0.0001 | 1,10 | 1 |
| *Danaus chrysippus* | Temperature | 1.234 | 12.424 | 0.293 | 1,10 | 0 |
| *Idea leucone* | Temperature | 0.000 | 176.991 | <0.0001 | 1,17 | 1 |
| *Cethosia cyane* | Temperature | 0.000 | 45.831 | 0.248 | 1,15 | 0 |
| *Vindula dejone* | Temperature | 10.199 | 289.120 | 0.005 | 1,17 | 1 |
| *Junonia almana* | Temperature | 5.830 | 24.134 | 0.034 | 1,25 | 1 |
| *Junonia atlites* | Temperature | 46.370 | 547.639 | <0.0001 | 1,11 | 1 |
| *Junonia iphita* | Temperature | 1.537 | 112.302 | 0.255 | 1,9 | 0 |
| *Doleschallia bisaltide* | Temperature | 31.481 | 192.753 | <0.0001 | 1,10 | 1 |
| *Bicyclus anynana* | Temperature | 34.304 | 185.891 | <0.0001 | 1,59 | 1 |

**Table S4:**
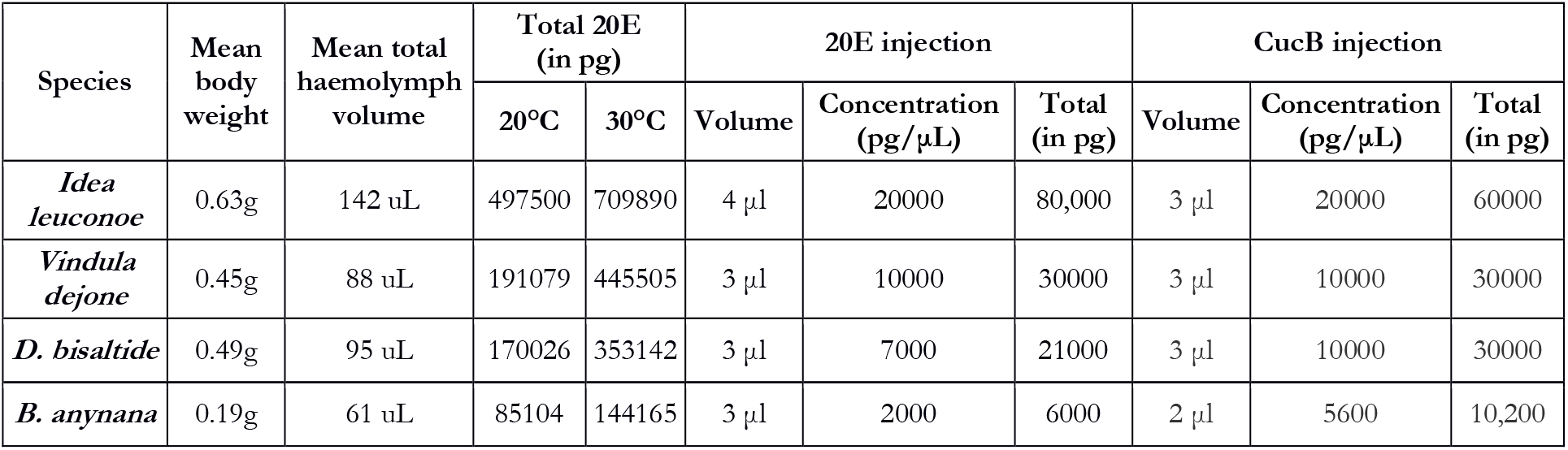
Mean body weight of wandering larvae, haemolymph volume and natural 20E titers at two different rearing temperatures; 20E and CucB injection volume. (N=5 for measurement of means)

#### Phylogenetic analysis

Patterns of plasticity in eyespot size were categorised in distinct groups based on positive, negative, or slopes undistinguishable from zero when eyespot size was plotted against temperature. Using a pruned version of a larger phylogenetic tree for all nymphalid genera^30, 31^, ancestral trait reconstructions were performed and evolution of the reaction norm slopes were mapped using maximum parsimony in Mesquite. Similar analyses were performed using data obtained for hormone titer plasticity where species were categorised into two categories – those with a positive slope or a zero slope, and data for presence or absence of EcR expression, and 20E-EcR signaling affecting eyespot size.

We also evaluated several hypotheses concerning the evolution of relevant traits with likelihood ratio tests (LRT) and Akaike Information Criteria (AIC). For all analyses, specific ancestral nodes of interest were “fixed” for a particular state and the resultant maximum likelihood score was used for LRT and AIC comparisons^18^. We performed four tests in all, investigating (1) whether the most recent common ancestor (MRCA) to all butterflies (node 14 in Fig. S1) had plasticity in spot and eyespot size or not; (2) whether the MRCA to all butterfly species with eyespots (node 17) had plasticity in eyespot size or not; (3) whether the MRCA to all butterflies (node 14) had positive hormone titre plasticity or not; and (4) whether the MRCA to all butterfly species with eyespots (node 17) expressed EcR in the locations of future spots / eyespots or not. For tests of eyespot size plasticity, we used a three-state coding scheme: positive size plasticity, negative size plasticity, and no plasticity. Character states were scored based on the sign of the slope of the reaction norm; species with reaction norms that were not significantly different from zero were scored as having no temperature-dependent plasticity in eyespot size (Table S2). Tests on positive hormone titre plasticty and EcR expression used characters coded as binary states. For AIC comparisons, we used the correction for small sample sizes (AICc) and evaluated models based on the AICc weight, *w_i_* = e^((min(AICc − AICc)/2)^. Models were considered significantly different if they differed by 2 or more log-likelihood units or the AICc weight was less than 0.2.

For all comparisons, there was little significant support for one hypothesis over another (Table S5). In tests on the origin of eyespot size plasticity, both the MRCA to all butterflies and the MRCA to all butterflies with eyespots had slightly better likelihood and AICc scores for being non-plastic than being plastic. Positive hormone titre plasticity in the MRCA to all butterflies had more support than a non-plastic MRCA, although the difference in likelihoods and AICc was not significant. Finally, the absence of EcR expression in the MRCA of all eyespot-bearing butterflies had higher likelihood and AICc scores than a model in which the MRCA did express EcR in future spot / eyespot centers. The absence of significant support for one model over another is largely due to the low number of species examined.

**Table S5:** Results of likelihood ratio tests and AIC comparisons. See Fig. S1 for node identities.

| Character | Node | State | -lnL | $\Delta\ln L$ | AICc | $w_i$ |
| --- | --- | --- | --- | --- | --- | --- |
| Size plasticity | 14 | Negative slope | 14.291 | 0 | 30.917 | 1.0 |
|  |  | Flat slope | 14.559 | 0.267 | 31.450 | 0.766 |
|  |  | Positive slope | 15.084 | 0.792 | 32.501 | 0.453 |
|  | 18 | Negative slope | 14.195 | 0 | 30.724 | 1.0 |
|  |  | Flat slope | 14.948 | 0.453 | 31.630 | 0.636 |
|  |  | Positive slope | 15.120 | 0.925 | 32.574 | 0.397 |
| 20 Hormone titre | 14 | Positive plasticity | 7.285 | 0 | 19.661 | 1.0 |
|  |  | No plasticity | 7.716 | 0.431 | 20.523 | 0.650 |
| EcR expression | 18 | EcR absent | 9.112 | 0 | 20.558 | 1.0 |
|  |  | EcR present | 9.604 | 0.491 | 21.571 | 0.612 |

**Fig. S1.**
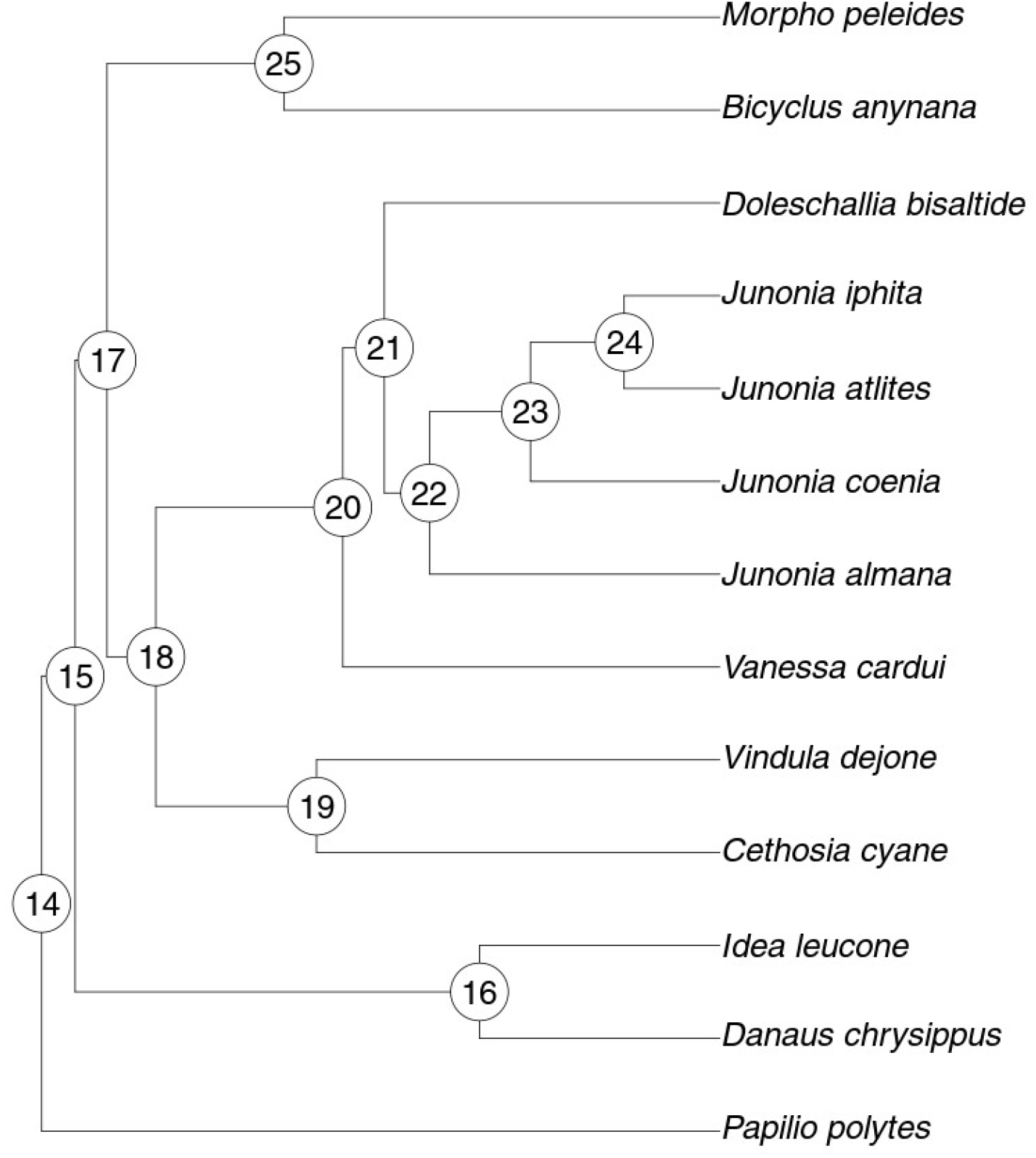
Tree used for ancestral state hypotheses tests. See text for explanation of node numbers.

**Fig. S2.**
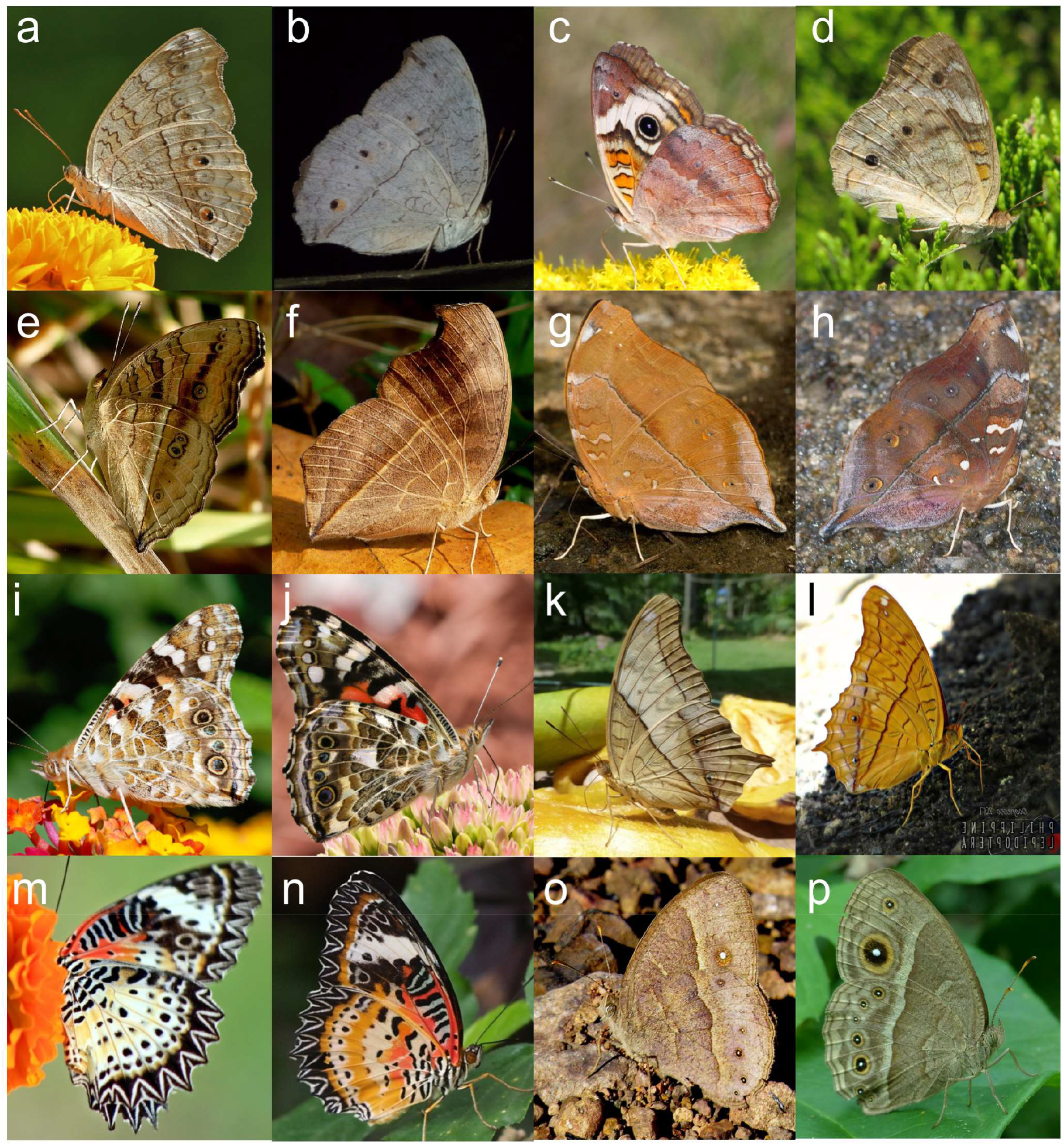
Phenotypic plasticity in wing patterns is observed across a wide variety of species in wild. a,b-Wet and Dry season forms of *Junonia atlites;* c,d-Seasonal forms of *Junonia coenia;* e,f-Seasonal forms in *Junonia alamana*; g,h-Differences in wing patterns across seasons in *Doleschalia bisaltide;* i,j-*Vanessa cardui* produces exquisite seasonal phenotypes; k,l-Seasonal variations in *Vindula dejone*; m,n-Seasonal forms in *Cethosia cynae*; o,p – Dry and Wet seasonal forms of *Bicyclus anynana*. Pictures are collected from crowdsourced repositories and copyrights belong to respective owners. Seasonal forms have been associated with reported time of collection.

## Acknowledgements

This work was supported by Singapore Ministry of Education award MOE2014-T2-1-146 to A.M. We thank Ms. Mei Lee Wong, Mr. Andy Loke and Mr. BT Chin (Penang Butterfly farm, Malaysia) for their support and supplies of butterflies used in these experiments. Work at SLING (M.R.W.) is supported by grants from the National University of Singapore via the Life Sciences Institute (LSI), the National Research Foundation (NRFI2015-05), and a BMRC-SERC joint grant (BMRC-SERC 112 148 0006) from the Agency for Science, Technology and Research (A*Star). We acknowledge Anne K Bendt for excellent SLING scientific program management and operations support. The EcR 10F1-s developed by Riddiford, L.M. was obtained from the Developmental Studies Hybridoma Bank, created by the NICHD of the NIH and maintained at The University of Iowa, Department of Biology, Iowa City, IA 52242.

## Author Contributions

Conceptualization: AM, SB

Methodology: SB, AM, MRW, JCO

Investigation: SB, LSHJ, FN

Formal analysis: SB, LSHJ, JCO, AM

Supervision, Funding Acquisition: AM

Writing – Original Draft Preparation: SB

Writing – Review and Editing: AM

